# Evaluation of *in vitro* and *in vivo* toxicity of pristine molybdenum disulphide nanosheets in Swiss albino mice

**DOI:** 10.1101/2021.05.07.443109

**Authors:** Umakant Yadav, Vimal Singh, Himanshu Mishra, Preeti S. Saxena, Anchal Srivastava

**Affiliations:** Department of Zoology, Institute of Science, Banaras Hindu University, Varanasi-221005, India; Department of Physics, Institute of Science, Banaras Hindu University, Varanasi-221005, India

**Author notes:** **Corresponding authors**, (Dr. Preeti S. Saxena).

**Keywords:** MoS_2_, oxidative stress, haematology, histopathology, toxicity, antioxidants

## Abstract

The Molybdenum disulfide nanosheets (MoS_2_-NSs) thin films has received increasing attention recently due to their versatile multi functionality including catalytic properties, photoluminescence and flexibility, which suggests their future, uses for biomedical applications. However, there are no studies in detail related with biocompatibility of MoS_2_ thin sheets. Here, weevaluated the dose-dependent effects of MoS_2_-NSs on cell viability (MTT assay) and release of lactate dehydrogenase (LDH) into culture media using MG-63 cells, as well as haemolysis, hematological, serum biochemical, antioxidants and histopathological parameters in *Swiss albino* mice. The MoS_2_-NSs was synthesized *via* facile hydrothermal method and characterized using XRD, Raman, SEM, TEM and HRTEM. The *in vitro* study results suggest that at lower concentration MoS_2_-NSs does not causes any toxicity. The lethal dose (LD50) was evaluated by intraperitoneal administration with different concentrations and estimated as ~1.0 mg kg^-1^. The higher dose (1.5 mg kg^-1^) of MoS_2_-NSs showed significant alteration in hematological markers and serum biochemical enzymes, as compared to control. Lipid peroxidation also shows significant alteration with respect to the control. Histopathological, hematological and biochemical examination, revealed no remarkable changes at lower concentration (less than 1.0 mg kg^-1^), however, higher concentration (1.5 mg kg^-1^) causes significant histopathological, antioxidants and biochemical alterations in tissues and serum, respectively. The results suggest that the lower concentration of MoS_2_-NSs can be used in future biomedical applications.

## 1. Introduction

In recent years, transition metal dichalcogenides (TMDs), a class of 2D nanomaterials, have spurred different areas of engineering, material science, and chemistry due to their unique physiochemical properties[1,2]. TMDs such as MoS_2_, WS_2_, MoSe_2_ and WSe_2_ are hexagonal layers and can be symbolized with the general formula MX_2_, where M represents a transition metal element from (Mo and W) and X represents a chalcogen (S, Se, Te). TMDs have layered structures arranged in the X-M-X conformation with in-plane covalent bonding and weak Van der Waal’s forces of attraction between layers[3,4].Molybdenum disulfide (MoS_2_), a member of TMDs family,has been studied extensively analogous to graphene[5–7].

In recent years, TMDs have attracted attention as a new non-viral gene delivery vehicle and in cancer therapy[8–10]. Reports are available that appropriate surface modification of molybdenum disulfide nanosheets (MoS_2_-NSs) with PEG and PEI served as a unique 2D nanocarrier for efficient siRNA delivery for PLK1 gene silencing[11], sensing applications[12], bone tissue engineering[13]and bioimaging[14]. The electrochemical sensing behavior of single-layer MoS_2_-NSs has been described, for glucose sensing as well as for the specific detection of dopamine[15,16]. In addition, MoS_2_-NSs have been also reported as template for hybridization of DNA strands and immunoglobulin’s for sensing of dopamine and ascorbic acid[17,18]. MoS_2_ act as guided missile for targeting of lung cancer. High mechanical strength of MoS_2_-NSs serves as an efficient reinforcing material for load bearing application and in tissue engineering[19].

Despite of its (MoS_2_) countless applications, there is no report available on MoS_2_-NSs *in vivo* toxicity in animal models. Therefore, as a successful candidate for *in vivo*applications in bio- and nanomedicine,*in vivo* study of MoS_2_-NSs is highly desirable.

Keeping above points in view, here we have evaluated*in vivo* toxicity studyofMoS_2_-NSs in *Swiss albino* mice in terms ofhematological, serum biochemical analysis, lipid peroxidation assay, antioxidants enzymes analysis and histopathological examinations.

## 2. Experimental Details

### 2.1 Chemicals

Sodium molybdatedehydrate (Na_2_MoO_4_.2H_2_O), hydrochloric acid (HCl), ethanol, hematoxylin–eosin stain and glacial acetic acid were purchased from Sigma Aldrich, New Delhi, India. Xylene, ethyl alcohol, paraffin wax, and microscope slides were obtained from Thermo-Fisher Scientific Pvt. Ltd. India. Serum biochemical enzymes (AST, ALT, and ALP) kits were procured from Beacon Pvt. Ltd. India. Antioxidant assay reagents were purchased from Merck Pvt. Ltd. India. All the chemicals and reagents were of analytical grade and used without any further purification.

### 2.2 Preparation of MoS_2_-NSs

MoS_2_-NSs have been synthesized using facile and eco-friendly hydrothermal methodwith some modifications[20]. In a typical synthesis route, 0.25g sodium molybdate and 0.5g of L-Cysteine were dissolved in 25 ml and 50 ml di-ionized (DI) water separately. These two solutions werestirred for 10 minutes at 40°C for complete dissolution of the precursors. Both the solutions were poured in a single beaker and mixed under constant stirring. The pH of the solution was maintained at ~5 using 0.1M HCl. Finally, the solution was transferred to a 100 ml capacity stainless steel lined Teflon autoclave. The Teflon autoclave was put in an oven and temperature of the oven was maintained at 230°Cfor 40 hour. After the completion of the reaction the black color sample has been taken out, which was washed with ethanol and distilled water thrice. The washed sample was dried for 12 hours at 60°C in open atmosphere.

### 2.3 Cell Culture

The MG-63 cells (human osteoblast) were procured from National Centre for Cell Sciences (NCCS)Pune, India and culturedin standard Dulbecco’s Modified Eagle Medium(DMEM), supplemented with 10% fetal bovine serum (FBS), and1% penicillin/streptomycin antibiotics under controlled atmospheric conditions(37°C temperature and 5% CO_2_). The old culture medium was replacedevery 2–3 days and cells were sub-cultured after 3–4 days whencell confluency reached about 80-85%.

### 2.4 Cell Viability

MTT (3-dimethylthiazol-2, 5-diphenyltetrazolium bromide) assay was perform to evaluate the cell viabilityafter 24 h of cell seeding in the presence and absence ofMoS_2_-NSs suspensions. Typically, MG-63 cells wereseeded in 96 well culture plates with a density 1×10^5^cells/well andincubated for 24 h for initial attachment. After 24 h, themedia was replaced with medium having variable concentrations(2, 5,10, 20, 40, 80, 100, 200,400 and 800 μgml^-1^) of nanocompositesuspensions (Stock concentration 1.0 mgml^-1^) and incubated for 24h. After incubation cells were washed twice with PBS and incubated again with media containing 0.5 mgml^-1^MTTfor 4 h at 37 °C under 5% CO2to form formazan crystals. Finally,MTT containing medium was removed and 100μL of dimethylsulfoxide (DMSO) was added followed by incubation on a rockingshaker for 20 min at 37 °C to dissolute the formazan crystals. Aftercomplete dissolution, absorbance of supernatant was recorded at 540nm using a microplate reader (BioTek, USA). The wells without MoS_2_-NSssuspensions were used as controls. The values were represented as themean±standard deviation (n= 3). Cell viability was calculated usingthe formula.

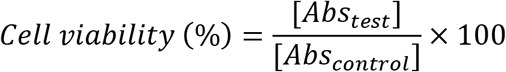

### 2.5 Lactate Dehydrogenase (LDH) release

Lactate dehydrogenase (LDH)assay was performed to evaluate the biocompatibility of MoS_2_-NSs.MG-63 cellswith a density 1×10^5^cells/wellwere seededin a 96-well plate and incubated with different concentrations(2, 5, 10, 20, 40, 80, 100, 200,400 and 800 μgml^-1^) of MoS_2_-NSs suspensions at37 °C under 5% CO_2_for 24 h.Standard LDH solution (1 ml) wasadded to the supernatant and the absorbance of thesupernatant was recorded at 490 nm to analyze the level ofLDH release.The wells without MoS_2_-NSs suspensions were used as negative controls and cells treated with Triton X-100 taken as positive control.

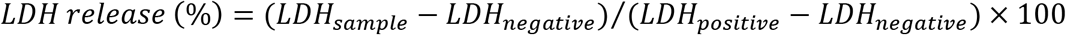

### 2.6 Haemolytic analysis

Haemolytic activities of MoS_2_-NSs were evaluated by detectingthe haemoglobin release from red blood cells (RBCs). Blood was collected in polypropylene tubes with anti-coagulant EDTA from healthy male *swiss albino* mice.The RBC was isolated from blood by centrifugation(3000 rpm at 4 °C for 10 min) and washed five times with PBSsolution (pH 7.4). Then, the RBC was diluted ten times with PBS. Furthermore, the RBCsuspension was added to theMoS_2_-NSssuspension in a volume ratio of 1:4 to give a final concentration of 25-800 μgml^−1^.PBS solution and Triton X-100were used as the negative (0% lysis) and positive (100% lysis)controls, respectively. The RBC suspensions were incubated at37 °C for 45 min. Subsequently, all the suspensions werecentrifuged at 3000 rpm at room temperature for 10 min, and the absorbance at 541 nm was recordedmicroplate reader (BioTek).The percentage haemolysis was expressed by the formula givenbelow.

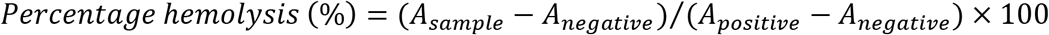

where, A_sample_, A_negative_, and A_positive_represent the absorbanceof the samples and negative and positive controls, respectively.

### 2.7 Animal Maintenance

All experiments with animal were carried out in compliance with the institutional ethics committee regulations and guidelines on animal welfare (Animal Care and Use Program Guidelines of IMS, BHU) and approved by Government of India.

Male *Swiss Albino* mice (~32 g), 3-5 weeks old, were procured from Institute of Medical Sciences, Banaras Hindu University, India. Mice were kept in cages (five mice per cage) and maintained under 12:12 hour light and dark photo period. Mice were fed with controlled *ad libitum* and water.

### 2.8 Lethal dose (LD50) estimation

For toxicological evaluation of MoS_2_-NSs, LD50 was evaluated using Irwin’s test method[21]. After acclimatization, twenty five mice were randomly divided into five groups, and each group contains five mice. One group was selected as control (0.9% saline), and rest four groups were selected as the experimental groups (0.5, 1.0 and 1.5 mgkg^-1^). After single dose administration of MoS_2_-NSs, animals were observed for behavior and clinical signs of lethargy for 8 days.

### 2.9 Study design

The homogeneous solution of MoS_2_-NSs was prepared in sterile 0.9% saline (Misonix, Long Island, NY, USA) using sonication at 40% amplitude with pulse 2 seconds on-off cyclefor 30 min. The concentration of the stock solution was 1.0 mgml^-1^. After acclimatization, twenty mice were randomly divided into four groups, each containing five mice. One group was selected as control (0.9% saline), and rest three as the experimental groups. The MoS_2_-NSs suspension was administered intraperitoneal to the experimental groups (I, II & III)having concentrations ~0.5 mg kg^-1^, 1.0 mg kg^-1^ and 1.5 mgkg^-1^,respectively. The time interval between dose administrationswas 24 h,up to 8 days. Body weight of eachmousewasrecorded before and after dose administration afterevery two days.

Different vital organs such as (liver, kidney and spleen) were excisedaseptically and weighted carefully for organ indices calculations. A portion of each organ (liver, kidney and spleen) were cut off and immediately fixed into Bouin’s solution for histopathology. The remaining tissueswere stored at −80 °C for further biochemical assay.

### 2.10 Hematological and Serum biochemical analysis

The mice were anaesthetized using diethyl ether and decapitated on eight days after six hours from the last dose. The blood was collected for hematological assays. The whole blood (~2.0 ml) was collected in polypropylene tubes with anti-coagulant EDTA (Ethylene diamine tetra acetic acid). Simultaneously, the serum has been separated by centrifugation of clotted blood (~37°C for 30 min) at 4000 rpm for 10 minute from whole blood maintained at 4°C. The serum was carried out to measure the level of different enzymes *viz*. alanine aminotransferase (ALT), aspartate aminotransferase (AST) and alkaline phosphatase (ALP) using colorimetric assay kits procured from Beacon Pvt. Ltd. India.Analysis of blood element *viz.red* blood cell counts (RBCs), hemoglobin (Hb), white blood cells (WBCs), mean corpuscular hemoglobin concentration (MCHC), platelet count (PLT), mean corpuscular hemoglobin (MCH), mean corpuscular volume (MCV) and hematocrit (HCT) were carried out using hematological auto-analyzer (MS-9-3 France).

### 2.11 Preparation of Homogenates

The different organs (liver, kidney and spleen) were excised out on ice-cold physiological saline and cleaned properly. The tissues were used to prepare a 10% w/v homogenate separately in 0.1 M phosphate buffer (pH 7.4) containing 0.1mM EDTA using a motor driven Teflon-pestle homogenizer (Fischer-Scientific), and centrifuged at 12000 rpm for 5 min at 4°C to separate the supernatant. After centrifugation, supernatant was collected and stored at −80 °C for further assays. The protein concentration was measured byBradford methodusing bovine serum albumin as standard[22].

### 2.12 Lipid peroxidation assay

The level of lipid peroxidation (MDA) in different tissues were measured using thiobarbituric acid (TBA) described by Ohkawa*et. al*.[23].The level of MDA was measured in nmolemg^-1^ protein. The supernatant was mixed with 2.8 mM Butylated hydroxyl toluene (BHT), 8.1% SDS, 20% Glacial Acetic acid and 0.8% Thiobarbituric acid (TBA) following a boiling at 100 °C in water bath for 1 hrs. The reaction was instantaneously transferred into running water and vigorously shaken with n-butanol: pyridine (15:1). The mixture was centrifuged at 1500g for 10 minute and the absorbance spectra of the supernatant have been recorded using UV-Vis spectrophotometer at 534 nm.

### 2.13 Superoxide dismutase assay

A 10% homogenates of different tissues were processed for SOD activity assay as per the mentioned protocol[24].100 μl of the different tissues homogenate were mixed in a 1.4 ml reaction mixture containing 20 mM L-Methionine, 1%(v/v) Triton X-100, 10 mM Hydroxylamine hydrochloride and 50 mM EDTA. After that 80 μl of 50 μM Riboflavin was added and the mixture was illuminated under 20W white light for 10 min. The reaction was stopped by adding freshly prepared Greiss reagent and recorded the absorbance spectra at 543 nm.

### 2.14 Catalase activity assay

10% homogenates of different tissues were processed for the assay in a reaction mixture comprising 0.8 mM H_2_O_2_, PBS and Potassium dichromate in Glacial Acetic Acid. The reaction was stopped by heating it in a water bath for 10 min and the absorbance spectra has been measured at ~570 nm to monitor the decrement in H_2_O_2_content. The residual H_2_O_2_ concentration after depletion by enzyme present in different extract was measured by spectrometer at ~570 nm and depicted as Catalase Activity. The standard curve was calibrated with varying concentrations of 0.2 mM H_2_O_2_ in PBS[25].

### 2.15 Glutathione peroxidase assay

Different tissue homogenates were handled for protein estimation. 150 μg of protein was mixed in a reaction mixture containing 50 mM phosphate buffer pH 7, 1 mM EDTA, 1 mM sodium azide, 0.5 mM NADPH, 0.2 mM reduced glutathione (GSH) and 1 unit of glutathione reductase. The reaction was put at room temperature for 1 minute to equilibrate. The reaction was started by adding 0.1 mM H_2_O_2_ and the decrease in the absorbance of reaction mixture was recorded at ~340 nm for 3 minutes at every 30 seconds interval in the UV-Vis spectrophotometer. The glutathione peroxidase activity was expressed as nanomole of NADPH oxidized/mg protein/ minute according to the procedure reported byMantha*et al*. [26].

### 2.16 Histopathological Analysis

The organs (liver, kidney and spleen) were surgically excised out from mice under diethyl ether anesthesia. A small piece of the tissue was excised from whole organs and washed with ice-cold normal saline (0.9% NaCl) and 20mM EDTA to remove blood traces. Further, the tissues were cut into small pieces and fixed immediately in Bouin’s solution for next 24 h. After fixation, the tissues were transferred to 70% ethyl alcohol and stored until processed. The tissue specimens (liver, kidney, spleen) were processed, embedded in paraffin, sliced into 0.5μm thicknesses, mounted on glass microscope slide and stained with hematoxylin and eosin (H&E) for histological examination under optical microscope. At least 10 slides were selected for histopathological evaluation of each sample.

### 2.17 Apoptotic cells analysis

Apoptotic splenocytes were spotted via Acridine orange (AO) – Ethidium bromide (EtBr) double stain. AO stained both types of cells i.e. apoptotic cells and viable cells emit green fluorescence. Early apoptotic cells exhibits membrane blebbing and chromatin strengthening as bright green spots or fragments while late apoptotic cells indicates orange to red nuclei having compressed or fragmented chromatin. EtBr stained only dead cells and emits red fluorescence when intercalated by DNA. Cells were centrifuged at 1000g for 5 minute to settled and add 100 μl of AO (100 μgml^-1^) and EtBr (100 μgml^-1^) in 1:1 ratio to the cell pellet. After 2–5 minutes a drop of stained cells were taken on a glass slide and mounted with a cover slip and examined under fluorescence microscope (Leitz MPV3, Wetzlar, Hesse, Germany) centered at ~520 nm emission for a ~440 nm excitation at 920 X magnifications[27].

### 2.18 Statistical Analysis

Statistical analysis was performed using SPSS 16.0 software. All experiments were repeated three independent times and all data were expressed as means ± SEM (standard error mean). One-way analysis of variance (ANOVA) followed by Dunnett’s post hoc test with *p*-values less than 0.05 was considered as statistically significant.

## 3. Results and Discussion

Applications of nanoscale materials have received a tremendous concern in nano-biotechnology due to their unique physiochemical properties. Prolonged exposure of nanomaterials may cause adverse effect on human health and environment. Particularly emerging applications of transition metal chalcogenides with unique optoelectronic and catalytic properties such as MoS_2_ for future biomedical applications warrants detail biocompatibility analysis *in vivo*. There is numerous in vitro biocompatibility analyses of surface modified MoS_2_-NSs. To the best of our knowledge this is the first report about the *in vivo*toxicity of MoS_2_-NSs with*Swiss albino* mice. Our results revealed that MoS_2_-NSs (1.5 mgkg^-1^) suppresses the activity of anti-oxidants and causes vital organ injuries.

### 3.1 Structural/microstructural and morphological characterizations of MoS_2_-NSs

Schematic in figure 1 demonstrate the detail plan for synthesis and *in vivo* toxicity evaluation. Structural/microstructural and morphological characterizations of MoS_2_-NSs have been performed using X-ray diffraction (XRD), Raman, scanning electron microscopy (SEM),and high resolution transmission electron microscopy (HRTEM)**(Figure 2)**. Figure 2(a) shows the XRD pattern of MoS_2_-NSs. XRD peaks have been fitted for Gaussian function. All the three peaks have been indexed for the hexagonal phase of MoS_2_ with space groups P63/mmc (JCPDS Card no. 37-1492).Plane (00.2) appears at ~13.99° which is slightly smaller than the peak position for bulk MoS_2_. Similarly, XRD peak corresponding to the (10.0) appears at ~38.17°, slightly larger than the corresponding value for bulk MoS_2_. This confirms that there is an expansion along [001] direction and in-plane compression[28]. Further, synthesized sample is characterized using Raman spectroscopy. A typical Raman spectrum is shown in Figure 2(b), where two peaks E^1^_2g_ and A_1g_pronounces at ~383 cm^-1^ and 409 cm^-1^ is observed. These two peaks correspond to the vibrational mode of first order Raman active center in plane and out of plane phonon vibration of 2H-MoS_2_-NSs respectively (Figure 2(c)). The spacing (Δ) between the peak positions E^1^_2g_ and A_1g_is found to be ~26 cm^-1^ which confirms that the synthesized MoS_2_-NSs are multilayer in nature[29].Figure 2(d) shows the SEM micrographof MoS_2_-NSs. It reveals that the MoS_2_-NSs are of few microns in lateral dimensions. TEM micrograph of MoS_2_-NSs has been shown in Figure 2(e). TEM image shows that the MoS_2_-NSs have agglomerated thin flakes structure. HRTEM image (Figure 2(f)) shows that the MoS_2_-NSs are multilayered in nature which is consistent with our Raman results. Inset (Figure 2(f)) shows the Fast Fourier Transform (FFT) pattern and interlayer spacing of MoS_2_-NSs. FFT image reveals that the stacking of layers is along c-axis. Interlayer separation is found to be ~0.68 nm, which corresponds to the (00.2) plane of MoS_2_-NSs. The interlayer separation corresponding to the (00.2) plane for MoS_2_-NSs is found to be slightly larger than the bulk (~0.62 nm), which shows that MoS_2_-NSs are swallowed along c-axis[30,31].

**Figure 1:**
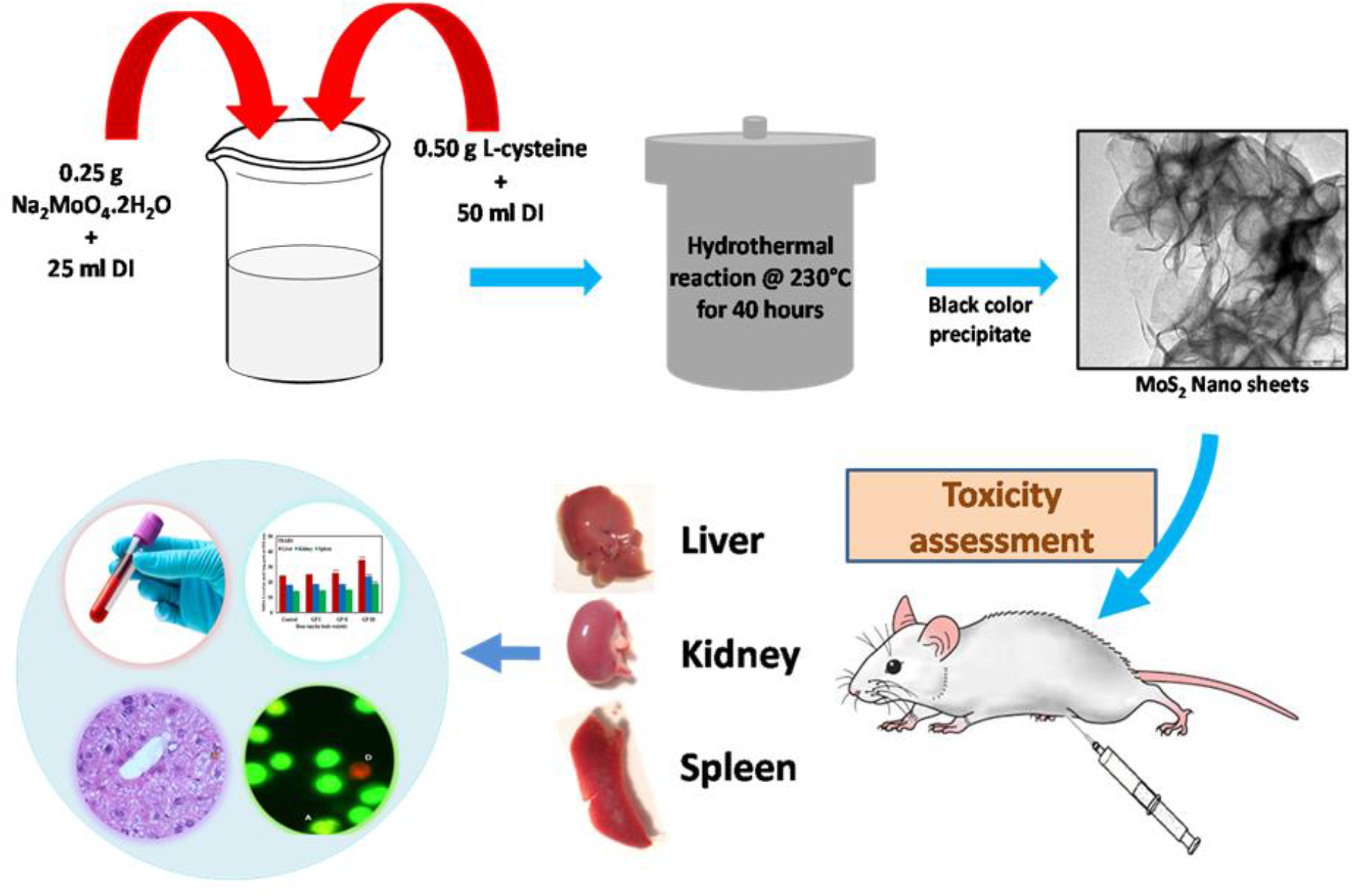
Schematic representation of for the synthesis and toxicity evaluation of MoS_2_-NSs.

**Figure 2:**
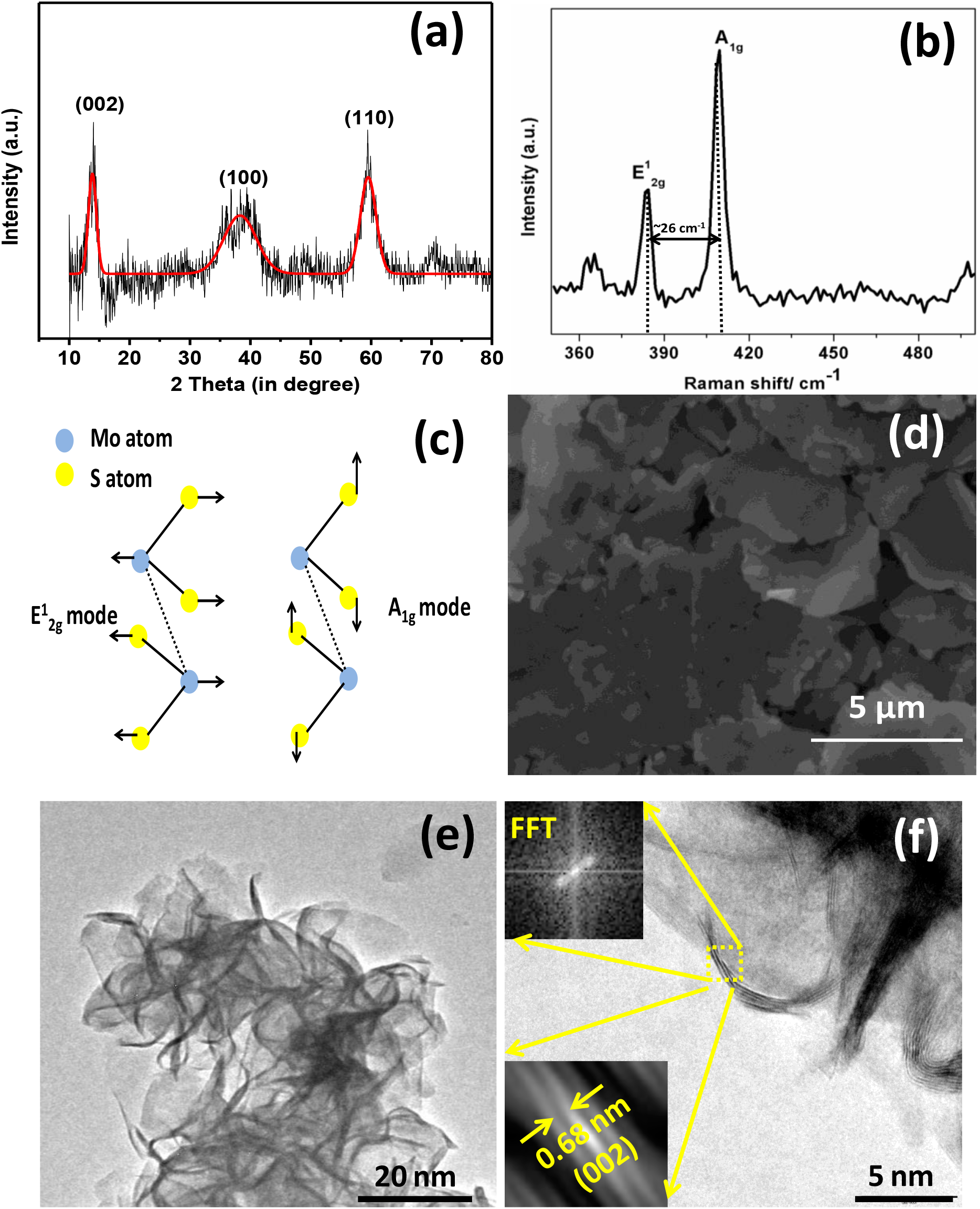
(a) XRD pattern, (b) Raman spectrum, (c) schematic for in-plane and out of plane vibrations, (d) SEM micrograph, (e) TEM micrograph, (f) HRTEM micrograph of multilayered MoS_2_-NSs; Inset shows the FFT pattern and interlayer separation of multilayered MoS_2_-NSs.

### 3.2 Cell Viability

Cell viability of the MoS_2_-NSssuspensions was investigated by MTT assay. The MG-63cells were incubated with different concentration of MoS_2_-NSssuspensions (0, 2, 5, 10, 20, 40, 80, 100, 200,400 and 800 μgml^-1^) for 24 h. Asshown in Figure 3a, results reveal that the viability of cellsincubated with different concentration of MoS_2_-NSssuspensions showed no significantdifference between them (p> 0.05) on 24 hup to 40 μgml^-1^. This shows thatthe cell viability was not affected by MoS_2_-NSs. Meanwhile, cellsincubated withhigher concentration than 80 μgml^-1^of MoS_2_NSs suspensions show a significantdecrease in cell numberFigure 3a. It may be due to lack of proper washing of MoS_2_-NSs. Wu *et al*. and Farshid*et al*. reported that MoS_2_-NSs have goodbiocompatibility with human cell lines[32,33].While cells incubated with 400 and 800 μgml^-1^of MoS_2_-NSssuspension shows significant decreased cell viability. Availablereports suggest that toxicity of MoS_2_-NSs depends on their synthesis routes[34]. Chang et al. have been reportedthat the level of toxicity of MoS_2_-NSs increases with theincrease of exfoliation[34].This is mainly due to the availabilityof large surface area and more active sites which may leads tothe ROS (reactive oxygen species) generation. Studies conductedby Zhang et al. for targeted imaging and photo thermal therapyusing RGD-QD-MoS_2_NSs against HeLa cells showed nosignificant toxicity[35].In our synthesis method, we have used sodium molybdate andL-cysteine as starting materials for the synthesis of MoS_2_-NSs. Inour case, some toxicity appears, it may due to two facts. First, if weconsider individual reagents, then hydrolysis of sodiummolybdate yields NaOH, H_2_O, and a small amount of MoO_3_that may be absorbed within the body or eliminated with urine.Further,L-cysteine is a semiessential amino acid and obtainedby the hydrolysis of animal materials, so it will also show notoxicity. Second, if we choose MoS_2_as an entity to discuss itstoxicity, then, as Yin et al. reported, the functionalization ofMoS_2_-NSs leads to improved antibacterial property. Ourhydrothermally synthesized MoS_2_-NSs is also functionalizedthat has been already in our earlier work[20,36].Thus, the results of cell viability demonstrate that MG-63cells have good proliferation rate in lower concentration of MoS_2_-NSssuspension.

**Figure 3:**
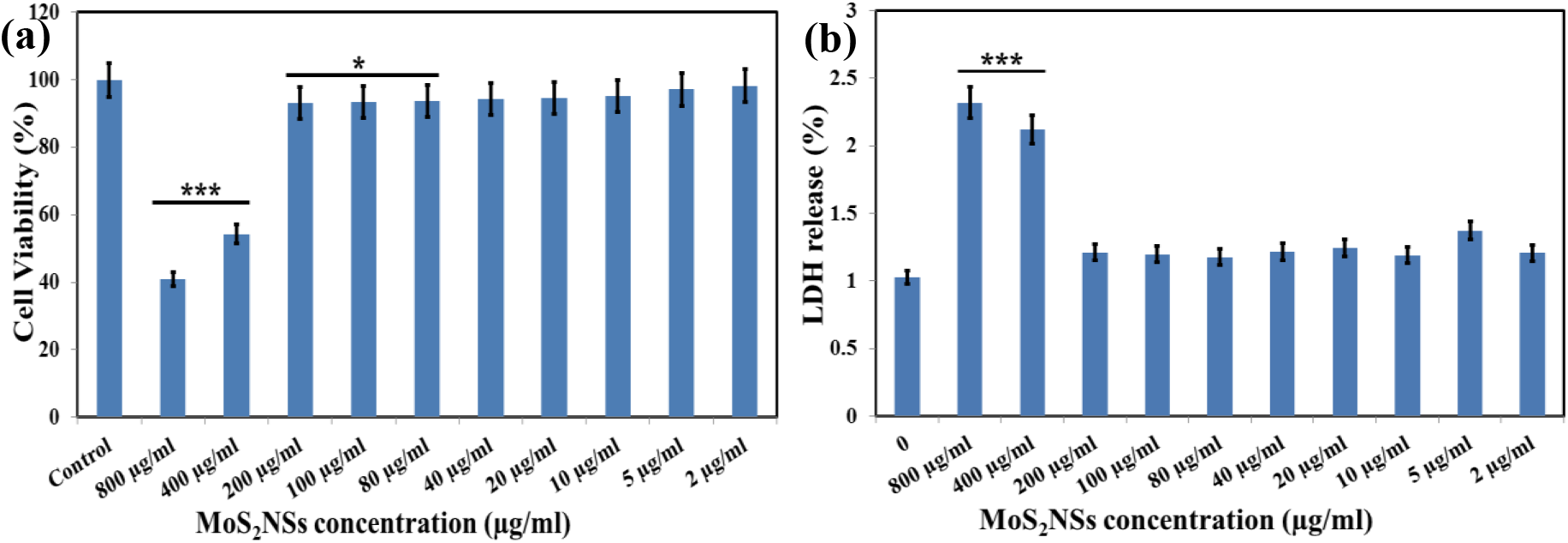
In vitro cytotoxicity tests: (a) MTT assay of MG-63 cells and (b) lactate dehydrogenase (LDH) release assay.Values are expressed as mean±standard deviation (*P< 0.05,**P< 0.01,***P< 0.001,n= 3).

### 3.3 LDH release

Lactate dehydrogenase(LDH) is a cytoplasmic enzyme that is found in all cells.In the case of mitochondrial and plasma membranedamage, intracellular LDH molecules are released into culturemedium. In our study, LDH release profiles were evaluated with respect toMoS_2_-NSsconcentration. From the result presented in Figure 3b, it was evident that cell toxicity and LDH release profiles do not show any potential toxicity, showing a lowLDH release amount (below 4%) and high cell viability aftertreatment. The results were in agreement with earlier reportsthat used PEG-GQDs for toxicity evaluation[37,38].

### 3.4 Haemolysis assay

The hemocompatibility study of any nanomaterial with blood components is a crucial toxicological parameter forthe application of nanomaterials for biomedicalapplications. To address the concern of blood compatibility, ahaemolysis assay was also performed against different concentrations ofMoS_2_-NSs (25–800 μgml^-1^) for 45 min and the result has been presented in Figure 4. The hemolysis percentages of MoS_2_-NSs were also calculated by taking OD at 541 nm. Theresult clearly shows that the RBCs incubated with 800 μgml^-1^of the MoS_2_-NSs, causes mild RBCs lysis.Therefore, our results demonstrated that MoS_2_-NSspossessed haemocompatibility in lower concentration[39].

**Figure 4:**
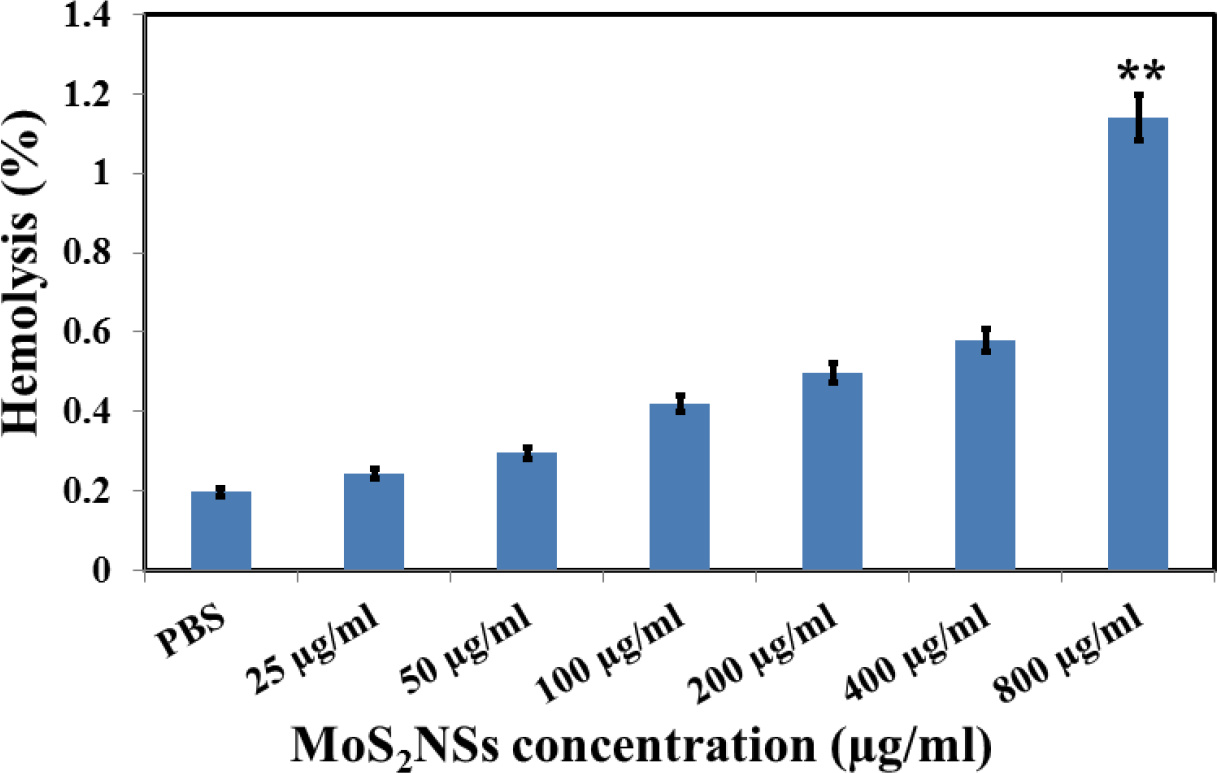
shows the percentage hemolytic behavior of MoS_2_-NSs.

### 3.5 Behavioral analysis and clinical parameters for LD50 determination

Based on the results of lethal dose (LD50), further doses of MoS_2_-NSs 0.5, 1.0 and 1.5 mg kg^-1^ were selected. No abnormal behavior or abnormal clinical signs were observed in doses of 0.5 and 1.0mg kg^-1^, while 40% death and other clinical and behavioral changes such as loss of appetite, tremor, loss of body weight and passive behavior were observed in the group receiving 1.5 mg kg^-1^ of MoS_2_-NSs.These changes occur due to effect of MoS_2_ toxicity at LD50.

### 3.6 Body and Organ weights

The body weight of first two group mice wassimilaras compare to control groups, and no significant difference was observed, whereas, on the fourth day, group-III (1.5 mg kg^-1^) mice shows significant increased body weight in comparison to control group (Figure 5a). Also, the different vital organs such as (liver, kidney and spleen) from MoS_2_-NSs treated mice, showed no significant weight differences with control groups (Table 1).

**Figure 5:**
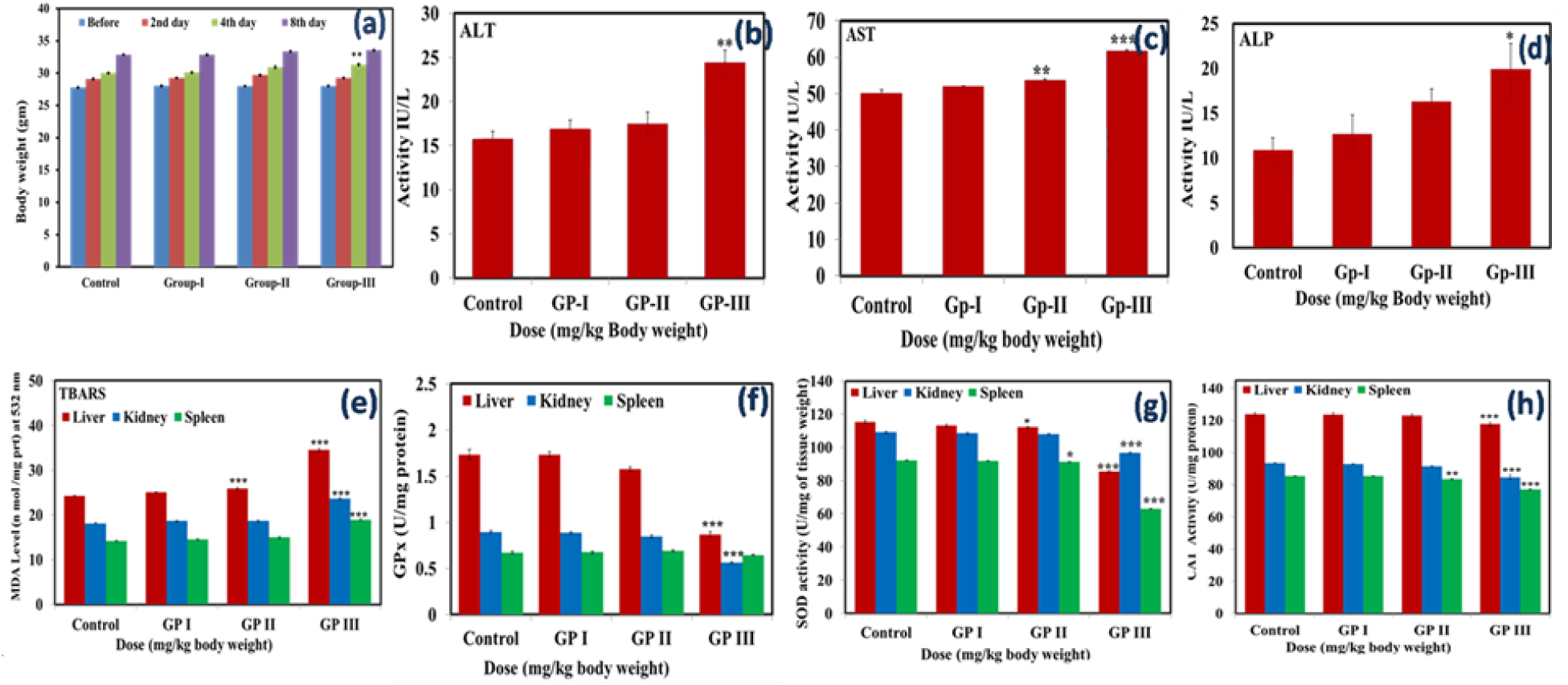
(**a**) Body weight of Control and MoS_2_-NSs treated mice. (**b**) Bar diagram shows the activity of Alanine aminotransferase (ALT/GPT). **(c)**Aspartate aminotransferase (AST/GOT) and **(d)**Alkaline phosphatases (ALP) in the serum of control and treated groups mice. Each experiment was done in triplicate. **(e)**Shows the level of TBARS in the homogenate of liver, kidney and spleen of treated groups and control. (**f)**the activity of SOD (**g)**shows the activity of catalase and (**h)**the activity of Glutathione Peroxidase. Each experiment was performed in triplicate. Data represents mean ±SEM; n=5 mice. At *P<0.05 statistically significant when compared to control.

**Table 1:** Vital organ indices of Control and MoS_2_-NSs treated mice at 8 day. Data represent (mean ± S.D.) (n=5).*P ≤ 0.05 vs. the control.(S.D. = Standard deviation)

| Organs | Control<br>(0.9% saline) | Group-I<br>(0.5 mg kg<sup>-1</sup>) | Group-II<br>(1.0 mg kg<sup>-1</sup>) | Group-III<br>(1.5 mg kg<sup>-1</sup>) |
| --- | --- | --- | --- | --- |
| Liver (gm) | 1.25 ± 0.11 | 1.27 ± 0.23 | 1.29 ± 0.13 | 1.43 ± 0.27 |
| Kidney (mg) | 212.2 ± 10.96 | 227.2 ± 11.78 | 213.6 ± 11.55 | 232.8 ± 27.67 |
| Spleen (mg) | 85.8 ± 6.81 | 83.6 ± 4.08 | 81.8 ± 4.54 | 85.6 ± 6.78 |

### 3.7 Hematological analysis

Major hematological markers such as red blood cell counts (RBCs), hemoglobin (Hb), white blood cells (WBCs), mean corpuscular hemoglobin concentration (MCHC), platelet count (PLT), mean corpuscular hemoglobin (MCH), mean corpuscular volume (MCV) and hematocrit (HCT) were evaluated.Here, significant increased levels of Hb and MCV were observed in group-II (1.0 mg kg^-1^) mice as compared to control. Furthermore, a significant decrease was observed in PLT count of group-III (1.5 mg kg^-1^) mice as compared to control (Table 2). These changes are observed due to interact of nanosheets with blood cells and causes various immunogenic response like inflammation and alteration of signaling pathways, which alters the hematological markers[40,41]. Detailed mechanism has been discussed in the forthcoming subsection.

**Table 2:** Hematological marker analysis of MoS_2_-NSs treated mice. Data represent (mean ± S.D.) (n=5). **P ≤ 0.05 vs. the control.

| Groups | Hb<br>(gm dl<sup>-1</sup>) | RBC<br>(10<sup>3</sup>UL<sup>-1</sup>) | WBC<br>(10<sup>3</sup>UL<sup>-1</sup>) | PLT<br>(10<sup>3</sup>UL<sup>-1</sup>) | HCT<br>(%) | MCHC<br>(%) | MCH<br>(pgm) | MCV<br>(fL) |
| --- | --- | --- | --- | --- | --- | --- | --- | --- |
| Control | 11.9 ± 0.43 | 8.00 ± 0.70 | 4.22 ± 0.64 | 5.04 ± 0.21 | 35.98 ± 1.02 | 33.2 ± 0.70 | 14.8 ± 0.44 | 44.2 ± 2.18 |
| Group-I | 11.78 ± 0.72 | 7.77 ± 0.78 | 4.32 ± 0.18 | 4.81 ± 1.33 | 36.38 ± 1.25 | 33.18 ± 0.75 | 15.2 ± 0.70 | 46.42 ± 2.04 |
| Group-II | 13.24 ± 0.90* | 8.00 ± 0.49 | 4.36 ± 0.21 | 4.62 ± 0.67 | 36.84 ± 0.81 | 33.78 ± 1.57 | 16.02 ± 0.61* | 48.98 ± 3.69 |
| Group-III | 12.66 ± 0.56 | 8.04 ± 0.37 | 4.32 ± 0.26 | 3.42 ± 0.53** | 37.94 ± 2.09 | 34.92 ± 1.24 | 15.7 ± 0.65 | 47.32 ± 5.21 |

### 3.8 Effects of MoS_2_-NSs on serum biochemical parameters

The blood serum of MoS_2_-NSs treated mice were analyzed for biochemical markers such as alanine aminotransferase (ALT), aspartate aminotransferase (AST) and Alkaline Phosphatase (ALP). The serum biochemical assays shown an increase in the alanine (ALT/GPT) activity for group-III (1.5 mgkg^-1^) mice. However, the highest doses of 1.5 mg kg^-1^ have found statistically significant effect in the increasing activity of ALT/GPT as compared with the control groups (Figure 5 b). Further the analysis of aspartate aminotransferases (AST/GOT) have been found a dose dependent increment on exposure to MoS_2_-NSs treated mice. However, the increase was less significant up to group II (1.0 mg kg^-1^) mice and becomes highly significant in group III (1.5 mg kg^-1^) mice as compare to control (Figure 5 c). Another enzyme, alkaline phosphatase (ALP), has shown an increased activity at highest concentration (1.5 mg kg^-1^) of MoS_2_-NSs treated mice compared with control groups (Figure 5 d).

The blood biochemical parameters were monitored to evaluate the function of liver, kidney and spleen after the administration of MoS_2_-NSs. The level of ALT has been tested along with AST and ALP to evaluate the liver function. It is seen that, whenever, liver become dysfunctional, the level of these blood biochemical enzymes (ALT, AST and ALP) rises.

According to our findings, significant increased level of ALT, AST and ALP (Figure 5 b, c and d) in serum might be associated with leak out of these enzymes in blood circulation due to triggered MoS_2_-NSs stress. Biologically, serum glutamic pyruvic transaminase (SGPT) is most suitable parameter for justification of liver injury than SGOT, because serum glutamic oxaloacetic transaminase (SGOT) is also found in kidney and heart muscles.Some previous studies have also reported that, in the case of hepatocellular injuries, the transportation of ALT and AST is disturbed which leads to leakage of ALT and ALP in general blood circulation and increase its activity[42,43].

On the other hand, the level of ALP in serum are associated to the function of hepatocytes[6]. In present study, we observed an elevation in the level of serum ALP (Figure 5(b)) in MoS_2_-NSs treated mice. For this elevation, we assume that MoS_2_-NSs must have affected the liver and may trigger the synthesis of ALP due to biliary pressure.

### 3.9 Antioxidant Enzyme Assay in Organs

Oxidative stress is considered as an important mechanism for the toxicity evaluation of nanomaterials. In this study, anti-oxidative indicators of the liver, kidney and spleen were evaluated. The results have been shown in Figure 5.The lipid peroxidation level, measured by the thiobarbituric acid reactive substances (TBARS) assay (Figure 5 e), shows a significant increment in the MoS_2_-NSs treated groups-II (1.0 mg kg^-1^) and group-III (1.5 mg kg^-1^) as compared with the control group (p < 0.05). The superoxide dismutase activity (SOD) (Figure 5 g) and the catalase activity (CAT) have shown a significant decrease in group-III (1.5 mgkg^-1^) as compared with control group (p < 0.05) (Figure 5 h). Likewise, the activities of glutathione peroxidase (GPx) were decreases significantly in group-III (1.5 mg kg^-1^) as compared to control group (p < 0.05) (Figure 5 f).

Oxidative stress is also an important parameter that playsa vital role in causing *in vivo*toxicity.To evaluate, the role of oxidative stress due to MoS_2_-NSs treatment, the level of Malondialdehyde (MDA) in the liver, kidney and spleen homogenates were assessed.The TBARS assay results revealed that the level of MDA increased significantly in the homogenates of liver, kidney and spleen of group-III (1.5 mgkg^-1^) treated mice (Figure 5 e), while a significant elevation in MDA level in liver homogenate only was reported in case of group-II (1.0 mg kg^-1^) treated mice. The elevation in the level of MDA primarily suggests that increased lipid peroxidation during tissue damage, suppresses the activity of anti-oxidants which prevent excessive free radical formation. In general, liver is the main organ for detoxification of hazardous materials and chemicals, So there is the high risk for reactive oxygen species (free radical) attack, leading lipid peroxidation and causes hepatotoxicity[44].Numerous other studies also supports lipid peroxidation *via* free radical generation due to nanoparticles induced toxicity[45,46]. Our results are also in accordance with other report that MoS_2_ increased lipid peroxidation in mice[47].

The result obtained from the present study clearly shows that the high dose (1.5 mg kg^-1^) administered in group –III mice has significant reduction in the level of SOD, CAT and GPx activity (Figure 5f,g,h) due to excessive production of reactive oxygen species (ROS).Li *et al*.reported that SOD activity could be suppressed due to excessive superoxide radicals formation and H2O2 accumulation[48]. The toxicity due to the higher dose of MoS_2_-NSs is mainly due to the generation of ROS. The detailed mechanism has been discussed in the forthcoming subsection.

Our results are also supported by other *in vivo* studies using different nanomaterials that possess the ability to elevate the oxidative stress. Similar to our findings, the experimental results using TiO_2_ has been confirmed to increase the hepatic injuries and significant alteration in antioxidants level[49,50].

### 3.10 Histopathological examinations

The histological sections of liver, kidney and spleen were observed. The morphological changes in the processed organs were examined. Figures 6 summarize the histological alteration in each group.

The microscopic observations of the control liver section, (Figure 6 a), showed normal structure and compact arrangement of hepatocytes. No obvious hepatic damage was observed in group-I and group-II in comparison to control group. However, significant morphological alteration like hepatocytes degeneration, mild necrosis, partial damage of central vein and hepatocyte vacuolation were observed in liver tissue (Figure 6 d) of group-III mice as compared to control group.

**Figure 6:**
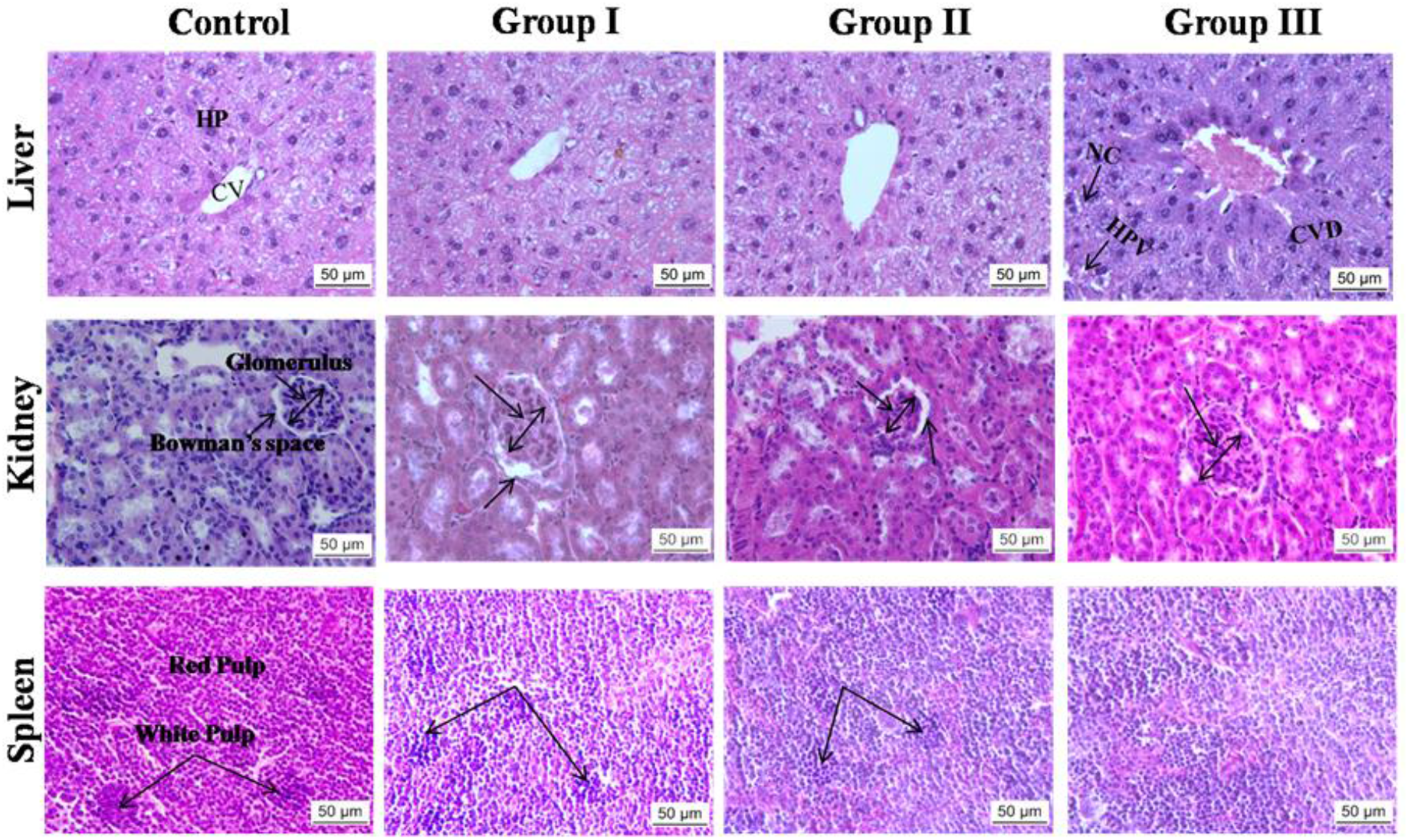
Histopathological evaluation (H&E staining, 40X) of organs in Swiss albino mice exposed to MoS_2_-NSs. (A,E,I) Control group; (B,F,J) Group I-0.5mgkg^-1^; (C,G,K) Group II-1.0 mgkg^-1^; (D,H,L) = 1.5 mg kg^-1^. (CV = central vein, HP=Hepatocytes, CVD= central vein damage, HPV = hepatocellular vacuolation NC = necrosis Glomerulus, Bowman’s space, white Pulp and Red Pulp).

Histopathological study of kidney shows no remarkable alteration in group-I and group-II mice, as compared with the control group. However, group-III (1.5 mg kg^-1^) mice (Figure 6h) show mild nephrotoxicity, such as swelling of glomerulus and decreased bowman’s space.

In case of spleen histological sections, none of the MoS_2_-NSs treated groups shows any significant pathological alteration, as compared with the control group (Figure 6 J,K,L).Further, histopathological evaluation was performed to observe any alteration in the morphology of the liver, kidney and spleen tissues. The histopathologies of liver of group-I (0.5 mg kg^-1^) and group-II (1.0 mg kg^-1^) (Figure 6 b, c) shows no morphological alteration in hepatocytes. However group-III (1.5 mg kg^-1^) mice show hydropic degeneration, mild necrosis, partial damage of central vein and hepatocyte vacuolation (Figure 6 d).

Kupffer cells are local macrophages of liver and involve in defense of liver against hazardous chemical and materials. Activation of these cells by hazardous chemical and materials induce to release inflammatory stuffs, growth modulators and ROS. Thus activation of Kupffer cells temper hepatic injury and chronic liver reactions[51]. Zhao *et. al*. reported that abdominal administration of nanoanatase TiO_2_ in mice causes kidney toxicity due to excessive generation of ROS, which leads lipid peroxidation and failure of antioxidant defense mechanism[52].

Histopathological study of kidney shows no remarkable alteration in group-I (0.5 mg kg^−1^) and group-II (1.0 mg kg^-1^) (figure 6 f, g) mice, as compared with the control group. However, group-III (1.5 mg kg^-1^) (Figure 6 h) mice show mild nephrotoxicity, swelling in renal glomeruli.

The results obtained from our study showed that high concentration of MoS_2_ nanosheets show abnormal signs. As an evidence, after autopsy we found that a large number of nanostructures are accumulated in intraperitoneal cavity of group-III (1.5 mg kg^-1^) mice and some are adhere onto the organs such as liver, kidney and spleen. Some previous studies also supports our result that nanoparticles can easily cross the wall of small intestine by absorption and then transported into different body parts like brain, lung, heart, kidney, liver, spleen and stomach through circulatory system[53].

### 3.11 Determination of apoptosis by Acridine orange–ethidium bromide (AO–EtBr) staining

The splenocytes cells isolated from the spleen of the MoS_2_-NSs treated mice showed an increased apoptotic bodies (Figure 7) as examined by the AO–EtBr staining. The dead cells that showed red color are few in number and the cause of death might be apoptotic as well as necrotic, however, we obtain most of the cells are viable emitting green colordue to acridine orange dye uptake.

The result obtained from AO-EtBr staining, clearly suggest that ROS must change the membrane potential of the spleenocytes which is the major reason for apoptosis of spleenocytes as shown by the AO–EtBr staining[27].

**Figure 7:**
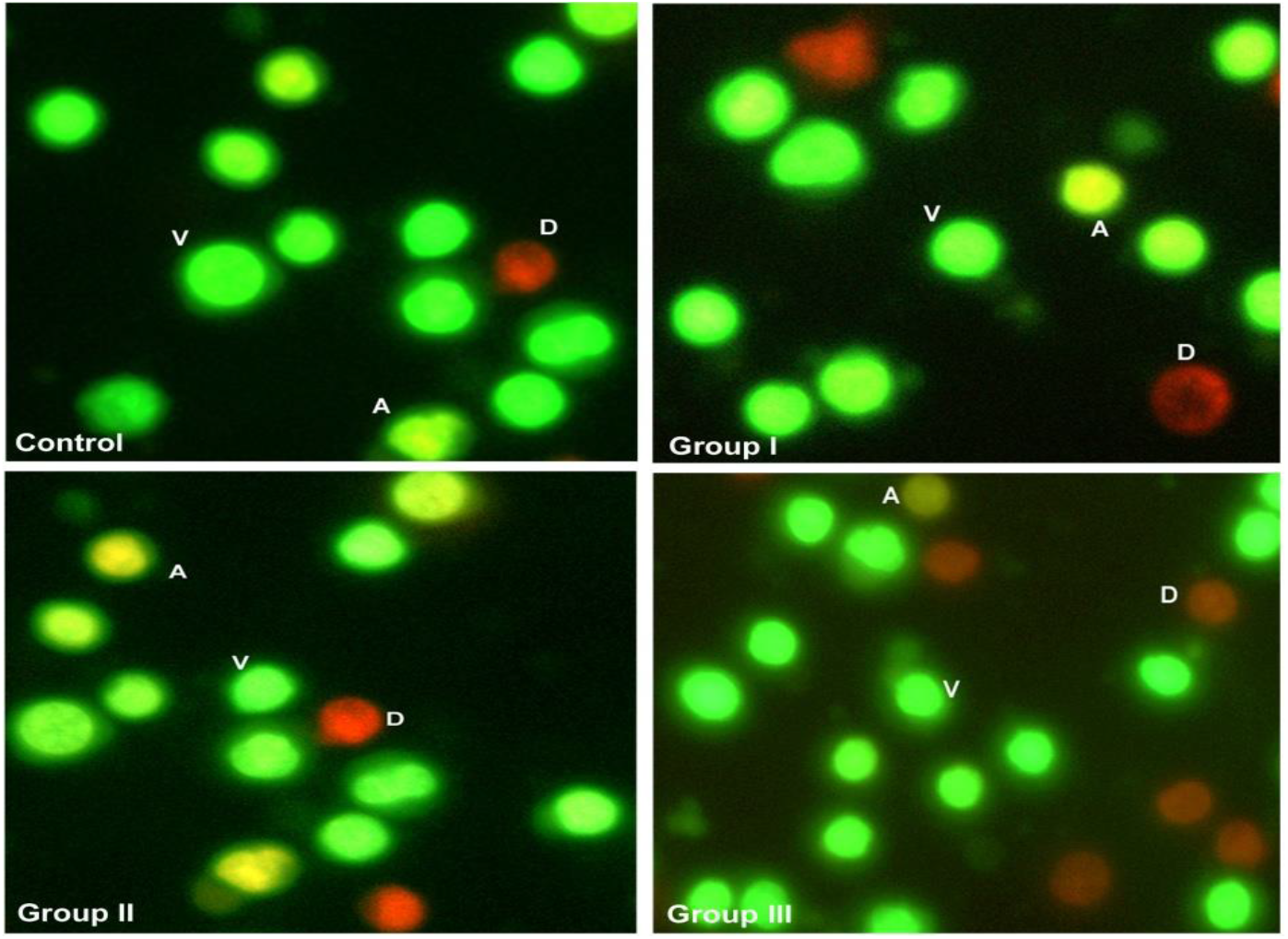
Staining of splenocytes with Acridine orange–ethidium bromide (AO–EtBr) showing apoptosis following the MoS_2_-NSs administration. Apoptotic cells (orange) are denoted by the letter “A” while the green are normal viable cells denoted by “V”. The red represents dead cells denoted by the letter “D”.

### 3.12 Mechanism for MoS_2_-NSs based toxicity

The mechanism of ROS generation by MoS_2_-NSs is shown in figure 8. Interaction of MoS_2_-NSs with Kupffer cell may activate it to produce inflammatory mediators, growth factors and reactive oxygen species (ROS)[51].Sulfur vacancies present at the edge of MoS_2_-NSs acts as electron donor to catalyze oxygen molecules in to superoxide ions[54]. These superoxide ionsare the precursor for most of the ROS generation[55].

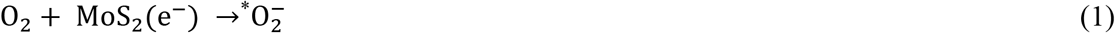

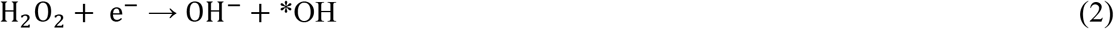

**Figure 8:**
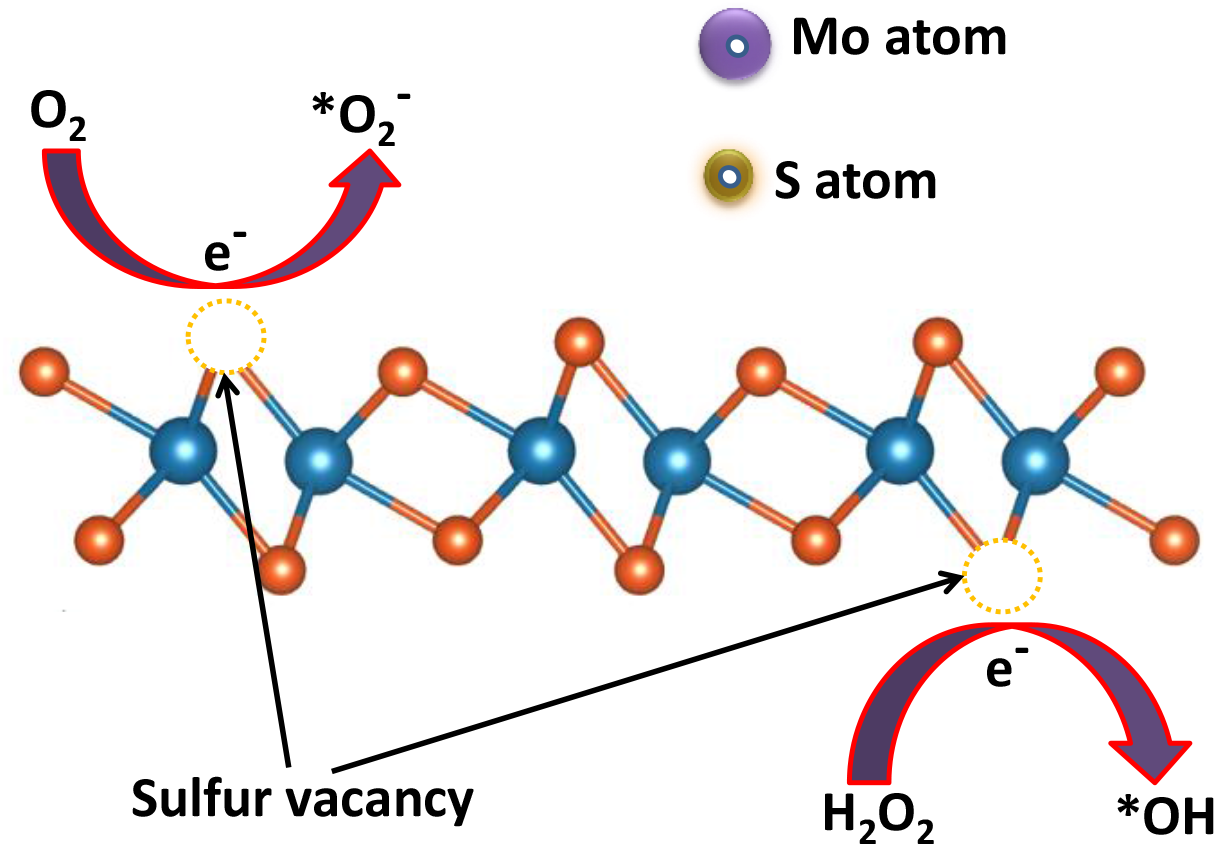
Schematic showing the mechanism of ROS generation due to MoS_2_-NSs.

These generated ROS may cause antioxidant depletion, hepatic injuries and increases the lipid peroxidation[56–58].

## 4. Conclusion

In summary, we have synthesizedMoS_2_-NSs by facile and eco-friendly hydrothermal methodusing sodium molybdate and L-Cysteine. Further*in vitro* and *in vivo*toxicity studies have been performed using differentconcentrationof MoS_2_-NSs. The *in vitro* study results suggest that at lower concentration MoS_2_-NSs does not causes any toxicity. While, *in vivo* study results conclude that serum biochemical markers were hiked significantly at higher concentration of MoS_2_-NSs (1.5 mg kg^-1^). Also, higher concentration significantly decreased the antioxidant enzymes activity. Histopathological studies in response to higher dose of MoS_2_-NSs treatment also shows altered architecture of liver as well as kidney, while spleen morphology remains unaltered. The toxicity of MoS_2_-NSs is supposed due to the ROS generation mainly. Due to significant importance of MoS_2_-NSs, present study suggests that lower concentration can be permissible concentration for the biomedical applications; however, improving the dispersion in aqueous media and surface coating of MoS_2_-NSsmight further allow application of higher doses. Furthermore, other toxicological parameters are also needed to evaluate the harmful biological responses associated with MoS_2_-NSs in future.

## Acknowledgements

Authors are thankful to UGC New Delhi for financially supported, CAS Department of Zoology and Physics, BHU, Varanasi, India. Authors are also thankful to DST, New Delhi, India for financial support under DST-PURSE (Scheme-5050).

